# Deciphering trophic interactions in a mid-Cambrian assemblage

**DOI:** 10.1101/2020.05.26.116848

**Authors:** Anshuman Swain, Matthew Devereux, William F Fagan

## Abstract

The Cambrian Period (541-485 Mya) represents a major stage in the development of metazoan-dominated assemblages with complex community structure and species interactions. Exceptionally preserved fossil sites have allowed specimen-based identification of putative trophic interactions to which network analyses have been applied. However, network analyses of the fossil record suffer from incomplete and indirect data, time averaging that obscures species coexistence, and biases in preservation. Here, we present a novel high-resolution fossil dataset from the Raymond Quarry (RQ) member of the mid-Cambrian Burgess Shale (7549 specimens, 61 taxa, ~510 Mya) affording new perspectives on these challenging issues. Further, we formulate a new measure of ‘preservation bias’ that aids identification of those assemblage subsets to which network analyses can be reliably applied. For sections with sufficiently low bias, abundance correlation network analyses predicted longitudinally consistent trophic and competitive interactions. Our correlation network analyses predicted previously postulated trophic interactions with 83.5% accuracy and demonstrated a shift from specialist interaction-dominated assemblages to ones dominated by generalist and competitive interactions. This approach provides a robust, taphonomically corrected framework to explore and predict in detail the existence and ecological character of putative interactions in fossil datasets, offering new windows on ancient food-webs.

**Significance Statement:** Understanding interactions in paleo-ecosystems has been a difficult task due to biases in collection and preservation of taxa, as well as low time resolution of data. In this work, we use network science tools and a fine scale dataset from the Cambrian period to explore: (i) preservation bias due to ecological/physical characteristics of taxa; (ii) evidence that the magnitude and sign of pairwise abundance correlations between two fossil taxa yields information concerning the ecological character about the interaction. All results in our work derive from using complex system approaches to analyze abundance data, without assuming any prior knowledge about species interactions – thereby providing a novel general framework to assess and explore fossil datasets.

## Introduction

Evolutionarily, the Cambrian Period (541-485 Mya) is unique because it witnessed the emergence and rapid diversification of phylum-level extant animal body plans and featured the highest morphological and genetic rates of animal evolution (Erwin et al., 2011; Lee et al., 2013). Morphological disparity and behavioral complexity increased (Carbone and Narbonne, 2014; Seilacher et al., 2005), prompting hypotheses about major shifts in ecological interactions and trophic structure during this period, due to major changes such as widespread predation and active (vertical) burrowing, which may have facilitated the first complex ‘modern’ food-webs (Dunne et al., 2008; Conway-Morris, 1986; Vannier and Chen, 2005, Erwin and Valentine, 2013; Mángano and Buatois, 2014). ‘Conservation lagerstätten’ sedimentary deposits, featuring exceptional fossil preservation of both ‘soft’ and ‘hard’ body features (Orr et al., 2003), permit detailed studies from which species interactions can be deduced (Butterfield, 2003).

Network-based studies provide critical insight on the structure and function of ecological systems (Delmas et al., 2018, Posiot et al., 2016; Ings et al., 2008), but paleo-assemblages often suffer from incomplete and indirect data (Roopnarine, 2010), time-averaging across large stratigraphic sections that obscure species coexistence (Kidwell and Bosence, 1991; Dunne et al., 2008; Roopnarine, 2010; Muscente et al., 2018), and biases in preservation, collection, and identification of both specimens and interactions (Koch, 1978; Smith, 2001; Dunne et al., 2008). Although some previous network studies have been performed on almost census preserved communities, such as in the Ediacaran (Mitchell et al., 2019; Mitchell and Butterfield, 2018; Muscente et al., 2019) Here, we report a new, extensive mid-Cambrian fossil abundance dataset featuring excellent preservation with high stratigraphic resolution, consistent taxa presence, and low biases in collection and identification. Using correlation network analyses of fossil abundance data and agent-based models, we find novel statistical evidence recapitulating 71 of 85 previously known or suspected species interactions, propose 117 previously unknown putative interactions, and identify a shift from assemblages dominated by specialist interactions to ones dominated by competition and generalized interactions. All results derive directly from fossil abundance data, without assuming any prior knowledge about species interactions.

Employing classic tools of network analysis in novel ways, we characterize fine-scale structure and dynamics of the paleo-ecological system represented by a novel 7549 specimen dataset from the Raymond Quarry (RQ) of the Middle Cambrian Burgess Shale of SE British Columbia, Canada (~510 Mya, Figure 1(a); S1). This dataset, which represents one of the most complete views of early animal assemblages, consists of species-wise abundance for 61 taxa in 10 cm levels across 9.3 m of shale. Previous network studies of paleontological data (e.g., Dunne et al., 2008) have focused on the Walcott Quarry, which is from the same geological period and has much higher species diversity than RQ, but unlike RQ lacks consistent species preservation across stratigraphic slices (Devereux, 2001). Therefore, this new dataset’s fine-scale spatial resolution and the site’s exceptional, consistent preservation of soft-bodied organisms offer advantages not available to most earlier paleo-assemblage network analyses (Dormann et al., 2017, Dunne et al., 2008). Moreover, in addition to unique usage of network methods, we utilized agent-based models to quantify and test key concerns, affording (a) a novel computational approach to understand preservation bias in fossil assemblages, (b) identification of putative interactions among taxa, (c) categorization of putative interactions into ecological roles, and (d) understanding of trophodynamics over time.

**Figure 1:**
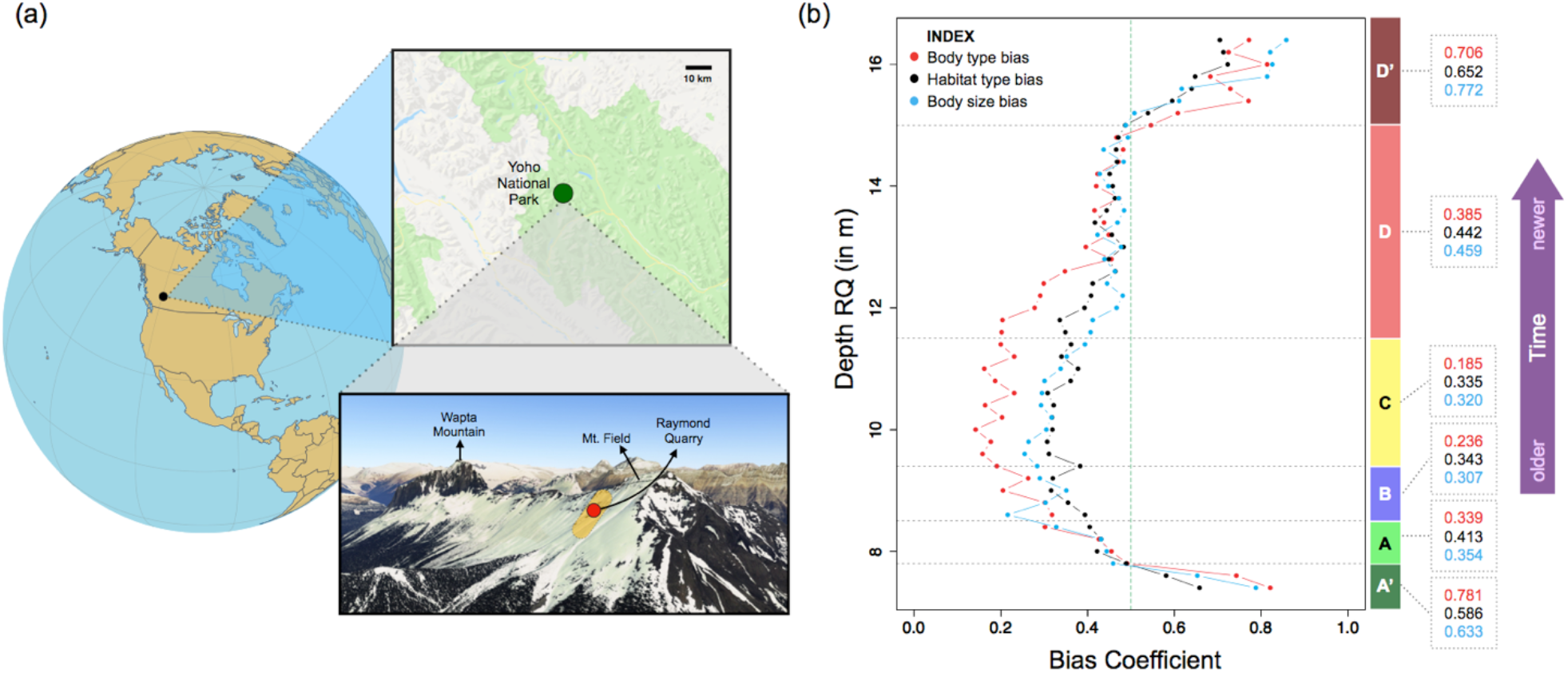
(a) Location of Raymond Quarry (RQ) (Yoho National Park, British Columbia, Canada), denoted by red dot; yellow region denotes extent of major Burgess Shale localities; Samples were collected from the RQ member along the ‘fossil ridge’ connecting Mt. Field and Wapta Mountain. (Figure S1, methods) (b) ‘Preservation bias coefficient’ for body type (with respect to soft, intermediate and hard bodied categorization) (in red), habitat type (in black) and body size categories (in blue) calculated using networks in running timeframe analysis, plotted along with estimated boundaries for the distinct sub-assemblages (A-D) in the 9.3 m shale section using variations of two different statistical approaches for biofacies detection: ANOSIM and SHEBI (Figure S2, methods) and the associated average bias coefficient for each sub-assemblage (in their respective index colors for each type of bias). The green dotted line depicts the bias threshold we adopted for inclusion versus exclusion of sample layers. Preservation bias coefficients exceeding 0.5 translate into substantial changes in the structure of interaction networks calculated from abundance correlations (Figure S3b). Note that the 10 cm layers comprising regions A’ and D’ were originally identified as belonging to sub-assemblages A and D, but fossils from A’ and D’ were not used in the analyses presented here because of evidence for high levels of preservation bias among taxa.

## Results and Discussion

Based on consensus results from two statistical approaches commonly used to define boundaries between fossil assemblages—SHEBI (Buzas and Hayek, 1998) and ANOSIM (Clark and Warwick, 1994) (methods, Fig 1(b); S2) —we identified four distinct sub-assemblages (named A-D in decreasing order of age), which match previous biofacies identification based on paleontologically defined trophic nuclei (Devereux, 2001). Based on these results, we calculated statistically corrected pairwise correlation of abundance for all taxa in each of the four sub-assemblages, and each of the 46 groupings of 20 cm levels organized to facilitate analyses at a finer depth scale (hereafter referred to as the running time-frame analysis) (methods). These correlations, with relevant regularization, were then used to construct correlation networks, for each sub-assemblage and each component of the running time-frame analysis. In these networks, each node was a taxon and each edge between a node pair represented significant correlation and thus possible interaction (methods).

Correlation networks can yield insights into possible interactions among taxa (Zhang, 2011; Carr et al., 2019; Barberán et al., 2012), but network features can be obscured by preservation and collection biases (Dunne et al. 2008; Carr et al., 2019; Jordano, 2016). Intensive sampling and detailed annotation reduced collection bias in this dataset, but preservation bias can still yield altered patterns of abundance. Statistical corrections have addressed some issues of fossil preservation biases (Flannery Sutherland et al., 2019; Starrfelt and Liow, 2016; Mitchell, 2015), but these have not targeted applications involving network analyses.

Preservation bias can occur for several reasons, most notably presence/absence of hard body parts/biomineralizable structures (which aid preservation), body size (which determines amount of preservable material and often the size of populations), and location/habitat (which provide differential conditions for preservation). If two taxa are both well preserved, their true abundance correlation is expected to exhibit less noise than correlations among pairs of taxa in which at least one taxon is not well-preserved. Differential preservation biases among taxa could introduce subtle structuring in a correlation-based network that would be biased towards more well-preserved taxa. The network-level consequences of such biases can be quantified by comparison with exponential random graph models (ERGMs), which have been used to understand the effects of bias and missing data (Robins et al., 2004).

To understand the influence of preservation bias on fossil correlation networks, we constructed an agent-based simulation model (ABM) of a complex resource-prey-predator system (consisting of 17 prey, 8 predators, and a common base resource for prey; methods). We ran 1 million simulations of this ABM that differed in various initial conditions, such as starting population size of each species and average resource density. For each of the 1 million simulations, we then created 100 cases, where each of the component species was assigned independently to one of three categories differentiated by probability of preservation (methods), and also retained the corresponding base case in which each species had perfect preservation (i.e., the original abundance data). For all 101 million cases (100 million cases with modified preservation and the corresponding 1 million original cases with perfect preservation), we then calculated abundance correlations among species pairs and constructed regularized correlation networks. We formulated a bias coefficient, using ERGMs and Hamming distance, to capture the effect of differential preservation on network structure through pairwise analyses of corresponding cases with modified and perfect preservation (Figure S3, methods). Higher bias coefficients corresponded to greater alterations of network structure.

We then applied this bias coefficient to the fossil data in the running time-frame analysis (methods). We separately considered three factors that could map onto differences in preservation: body type (hard-bodied, partially hard-bodied, soft-bodied), body size (<15cm, 15-30 cm, and > 30 cm maximum size), and habitat usage (endobenthic/epibenthic, nektobenthic and nektonic/pelagic) (methods). Information on these factors were compiled from literature surveys: body type, maximum body size, and habitat usage (see supplementary data).

For all three factors, we found evidence for significant differential preservation bias at the start and the end of the collected assemblage in regions A’ and D’ that were, respectively, originally part of sub-assemblages A and D identified through biofacies detection (Figure 1(b)). Because of their heightened preservation bias, which was strong enough to substantially alter apparent network structure, data from regions A’ and D’ were excluded from further analyses. In contrast, we found low levels of preservation bias coefficients in each of the defined sub-assemblages (A-D), with respect to body type, body size, and habitat (Figure 1(b); bias coefficient was < 0.5 for all cases; methods). These preservation biases were low enough to have inconsequential effects on network structure (Fig. 1b, Fig. S3). One would expect heightened levels of preservation bias at the beginning and end of a fossil bed if the strata above and below the sampled assemblage did not allow proper preservation of organisms due to a change in environmental (preservation) conditions (Orr et al., 2003). From a taphonomic viewpoint, factors such as differential transport experienced between taxa and between fossil beds, the degree of time averaging, and pre-burial transport distances may shape the preservation of discoverable assemblage contents as well (Olson et. al., 1980; Martin, 1999). Consequently, complete preservation of all ecological information is seldom expected (Flannery Sutherland et al., 2019; Saleh et. al., 2020). However, the consistency in low preservation bias coefficient with respect to body preservation type, habitat type, and body size throughout sub-assemblages A – D (with removal of A’ and D’ and corresponding running time frames) suggests that the net taphonomic effect resulted in an overall relatively homogeneous burial of a group of taxa across the whole assemblage (Figure 1(b)). Nevertheless, some loss of taxa may not have been inferable from the fossil data using our methods, and small differences in preservation may still be present throughout the assemblage at the taxon level.

Even though we report no significant preservation biases beyond those at the A’ and D’ ends of the RQ assemblage based on the predicted ‘in-situ’ preservation potential of the RQ taxa, the original interactions may still have been subjected to certain biases (Butterfield, 2003). However, these possibilities are not pertinently different than methodological biases affecting recent or extant ecological data (Dunne et al., 2008; Armitage and Jones, 2019).

Both experimental and theoretical studies predict that prey-predator abundances should be correlated on long time scales (Liebhold et al., 2004; Tobin and Bjørnstad, 2003; Blasius et. al., 2019), and we found this to be true for our ABM simulations (Figure S4). This result supports the premise that fossil abundance correlations might correspond to potential species interactions, where the degree of preservation bias is low, such as in extremely well-preserved assemblages like the Burgess shale (Saleh et al., 2020), and census-like preservation of Ediacaran communities (Mitchell et al., 2019). Furthermore, the distribution of abundance correlations should characterize system-level interactions. For example, we might hypothesize that abundances of competitors should be negatively correlated whereas abundances of species engaged in highly specialized interactions should be strongly positively correlated, assuming homogeneous transport and burial. The shape and location of the distribution of fossil abundance correlations differed among sub-assemblages A-D (Figure 2 (a), (b)). In particular, the frequency of small magnitude correlations and of negative correlations increased over time from A to D.

**Figure 2:**
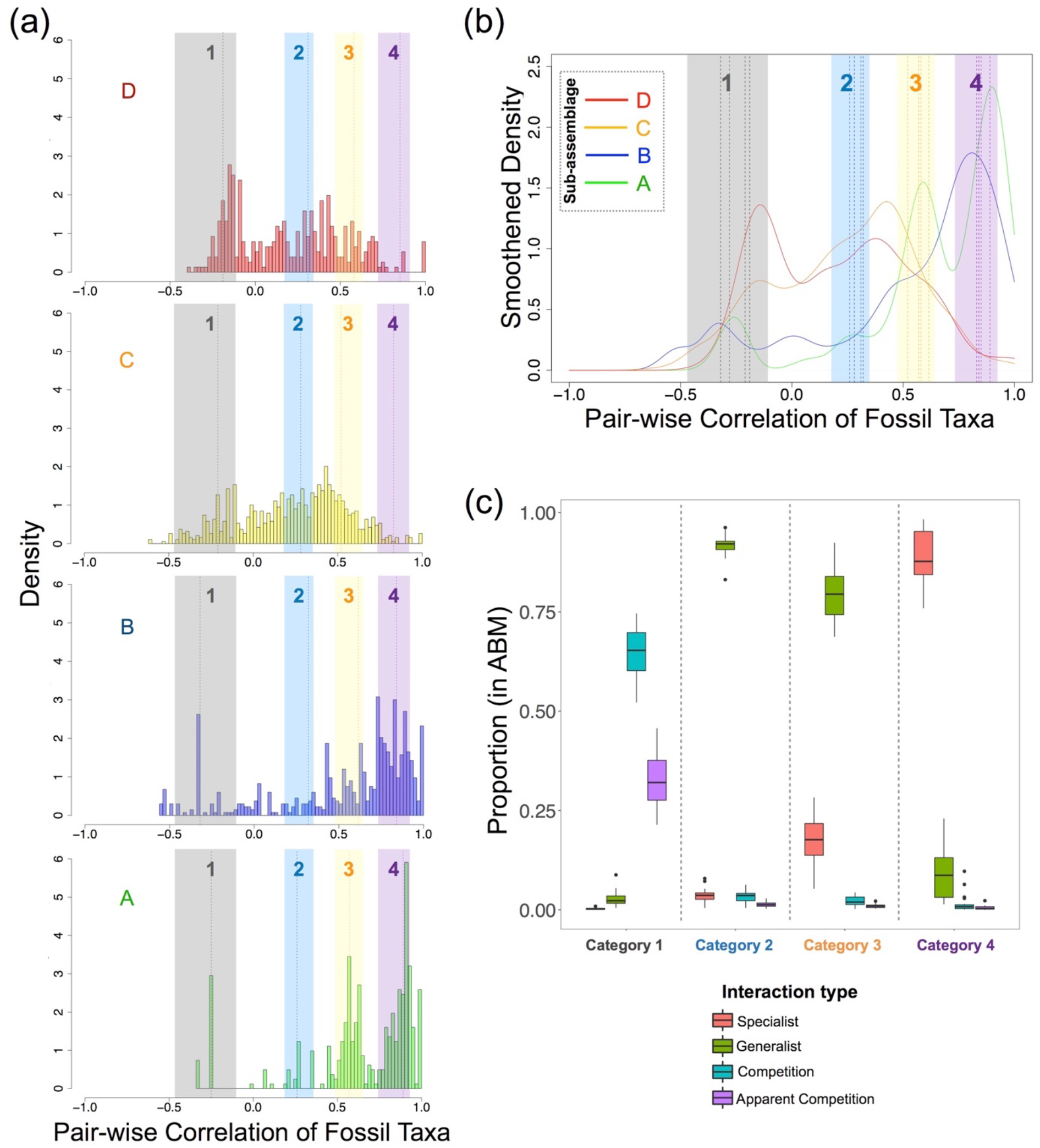
Characterization of interactions: (a) Distribution of pairwise correlations across A-D; dotted lines indicate the means of the basis Gaussians in each sub-assemblage calculated using maximum likelihood analysis for decomposing the abundance correlation distributions; colored bands indicate the four interaction categories 1-4, which represent the range of the Gaussian means for each interaction type, clustered from the basis Gaussians of the running timeframe analysis using spectral analysis with gap statistics (see methods). (b) Summary of the four categories of interactions, calculated from spectral analysis, on the smoothened density distributions of pairwise correlation of fossils in the four sub-assemblages A-D; dotted lines indicate the means of Gaussian basis distributions from A-D; (c) Proportion of different feeding types in the ABM whose abundance correlations fall in the colored bands identified in (a), suggesting, for example, that the negative correlations in category 1 (grey) are dominated by competition and apparent competition interactions whereas the strongly positive correlations in category 4 (purple) derive from specialized interactions.

To explore if changes in correlational distributions represented a shift in the dominant mode of species interaction over time, we decomposed the corrected correlation distribution for each sub-assemblage A – D, which were used in network construction, into its respective basis functions using maximum likelihood (methods). In each sub-assemblage A – D, the distribution of abundance correlations was best fit by a sum of four Gaussian distributions (Figure S6, methods), and across sub-assemblages, the four Gaussians had similar means but different amplitudes and variances (Figure 2 (a)). We found a similar result for the finer scale running timeframe analysis. To categorize these Gaussians in the correlations of fossil data into possible clusters, we calculated a pairwise similarity matrix of all the component Gaussians across the assemblage and then performed a spectral analysis with the ‘gap’ statistic (Tibshirani et al., 2000; methods) on it, which revealed four clusters of Gaussians (Figure 2 (a), (b); methods).

To understand the origins and potential meaning of the four clusters/categories of Gaussians in the empirical data, we used the ABM to simulate the dynamics of hypothetical ecological communities that differed in the importance of prey-predator interactions and competition. We considered trophic-relation-based ABM systems involving prey, specialist predators, and generalist predators, which implicitly also allow for competition and apparent competition (or, intra-guild competition). We compared where the abundance correlations associated with specialist, generalist, and competitive pairwise interactions from the simulated ABM communities fell relative to the four categories obtained via clustering from the fossil correlation distributions. Fully 86% of all correlations derived from interactions in the ABM fell within the intervals of the four empirically defined categories (Figure 2 (a), (b), (c)). In support of our initial ideas about the relationship between abundance correlation and interaction type, we found that ABM interactions involving competition and apparent competition dominate Category 1, generalist prey-predator interactions dominate Categories 2 and 3, and specialist prey-predator interactions dominate Category 4 (Figure 2 (c)). To explore, we used a second ABM in which each prey-predator interaction was weighted by the predator’s preference for that prey. From these simulations, we recovered the specialist-generalist spectrum of interactions, and further, identified a non-linear relation between a predator’s preferences for prey and the abundance correlations recovered from the ABM. The abundance correlation between a predator and its lower preference prey was weaker than that expected for the same prey unweighted by preference (Figure S4).

With reference to the ABM results, we can interpret that ‘Category 1’ (grey) involves negative correlations suggesting competition and apparent competition interactions among taxa (Fig. 2(c), leftmost column). Alternatively, such negative correlations can also arise if species have different habitat preferences and the relative availability of different habitats changes over time. Similarly, correlations in Categories 2 (blue) and 3 (light orange) likely involve generalist consumers. If a consumer is not specializing on one resource but eats many, it will be only loosely correlated with its prey (Fig. 2c, middle columns). A weak positive correlation could also mean that both members of a species pair use similar resources. Category 4 (purple) would derive from component Gaussians that feature consistently strong, positive correlations. This category likely represents specialist predation in which a predator depends solely or very strongly on a particular prey species (Fig. 2c, rightmost column). Alternatively, if two interacting species are strong mutualists or exhibit a strong joint dependence on environmental conditions, similarly strong positive correlations could emerge. Please note that throughout our work, we will refer to interactions as specialist or generalist (or competitive) without ascribing roles to particular taxa as we cannot ascertain which species are the prey and the predator in a given pair on the basis of abundance only. Therefore, our network is undirected.

We acknowledge that correlations may exist based on similar habitat/environment, and attempted to address this concern in three steps. First, we calculated the preservation bias for habitat (Figure 1(b)) and did not find any significant effect of habitat on the network structure. Next, to support our trophic ABM analysis as a benchmark for categorization, we tested whether two alternative reasons for correlations (i.e., habitat specializations for negative correlations, and habitat/environment sharing for positive correlations) impacted our analysis of the fossil data. To do this, we used a stochastic block model (SBM) on the sub-assemblages and on the running-time-frame data to see if the clustered distributions of interactions could be explained instead by habitat/environment and motility data hypothesized in literature (see Figure S10).

SBMs find community structure in networks (i.e., blocks of nodes which interact more among themselves than with others outside the block), and in this application would indicate clustering of interactions based on similar habitat or motility (Karrer and Newman, 2011). To implement the SBMs, we first computed the Shannon’s equitability index of each block in a given network and then calculated the weighted average of this index across blocks. An index value of 1 implies equal distribution of habitat or motility types across blocks, while a 0 indicates complete dominance of one type in a block. The SBM analyses revealed no strong dependence of the empirical correlations on either habitat or motility (Figure S10(c)).

Lastly, to explore whether negative correlations can arise from species specializing on different habitats, we compared the frequency of negative correlations within the same habitat, to the corresponding values for different habitats. For the running time scale data, there were no more negative interactions among taxa from different habitats than from the same habitats (Pearson’s *X*^2^ with Yates’ continuity correction: mean p-value (across the entire running time frame analysis) = 0.89, range = [0.68,0.93]), thereby excluding any strong effect of habitat exclusivity. We found similar results for positive correlations, again finding an absence of association between interactions and species’ habitat types (Pearson’s *X*^2^ with Yates’ continuity correction: mean p-value (for all time frame analysis) = 0.26, range = [0.19,0.35]). Collectively, these results suggest that habitat did not play a significant role in the structure, value and distribution of correlations in our network and that these correlations instead likely stem from species interactions.

Refocusing on the distributions of correlations, we observe that across fossil sub-assemblages A-D, negative interactions increase in relative frequency over time (Figure 2 (a), (b)). Similarly, specialist and generalist interactions increased from sub-assemblage A to B, but specialist interactions are largely absent in D, and occur only infrequently in C. Systematic change in the fossil transport regime could explain this, but this seems unlikely given the consistency of pairwise correlation signals over time (see fig S9). Alternatively, long term environmental change could have decreased regional productivity and made resources rare in the area of fossilization. Such long-term loss of productivity could have led to an increase in competitive interactions and loss of specialist interactions (including both mutualists and trophic interactions), which are more prone to extinctions (or extirpations) (Colles et al., 2009; Ryall and Fahrig, 2006). These results contrast with previous food-web based studies, where generalists dominate early structuring of food webs, followed by specialists (Piechnik et al., 2008). As such, instead of representing colonization of new habitats, our dataset may provide a window into fluctuating ecological abundances transported locally in a near-shore habitat, which were fossilized during intermittent episodes of exceptional preservation. Although we do not know the timeframe of deposition of RQ exactly—and it might be fairly long—there are no anatomical changes observable in the fossil taxa, which are consistently present throughout the assemblage. This suggests that the assemblage remains within an ecological regime rather than reflecting evolutionary time.

Analyses of abundance correlation networks recovered many proposed prey-predator interactions. We used the term ‘consensus interactions’ to refer to those species pairs whose abundance correlation yielded the same interaction categorization for more than 50% of the strata where the two taxa co-occurred. For species whose trophic interactions have been described or suggested in past literature and whose abundance in this dataset was sufficient for correlation analysis (Figure 3, innermost region), fully 83.5% (71 of 85) of consensus interactions predicted through our correlational analyses have been independently proposed in paleontological literature (Dunne et al., 2008; Erwin and Valentine, 2013; see methods). Our analyses supplement these expert propositions by assigning pairwise interactions into categories of prey specialization or preference by reference to correlation categories (Fig. 2). Furthermore, we propose 117 new possible pairwise trophic interactions based on abundance correlations identified here. These include 75 putative interactions for species whose trophic interactions were previously only partially known (Figure 3, innermost submatrix), and another 42 putative interactions involving species whose trophic interactions were not previously reported. Lastly, 18 pairwise interactions previously known from the literature could not be recovered here because the taxa involved were represented at very low densities in the fossil dataset (Fig. 3, submatrix with red background). Results in Fig. 3 only considered trophic interactions. Our correlational analyses also identified 137 possible competitive interactions for which there is no reference set because competitive interactions are more difficult to deduce from paleobiological data (Figure S7). Certain high correlation interactions may have been mutualisms, or based on shared environmental preference, common habitat patterns, or indirect interactions, rather than being trophic in nature (Freilich et. al., 2018). We searched, unsuccessfully, for a strong habitat dependence in the fossil data, but still cannot rule out any of these alternative possibilities with current data. Direct fossil evidence and further paleontological knowledge is needed to verify or explore these points.

**Figure 3:**
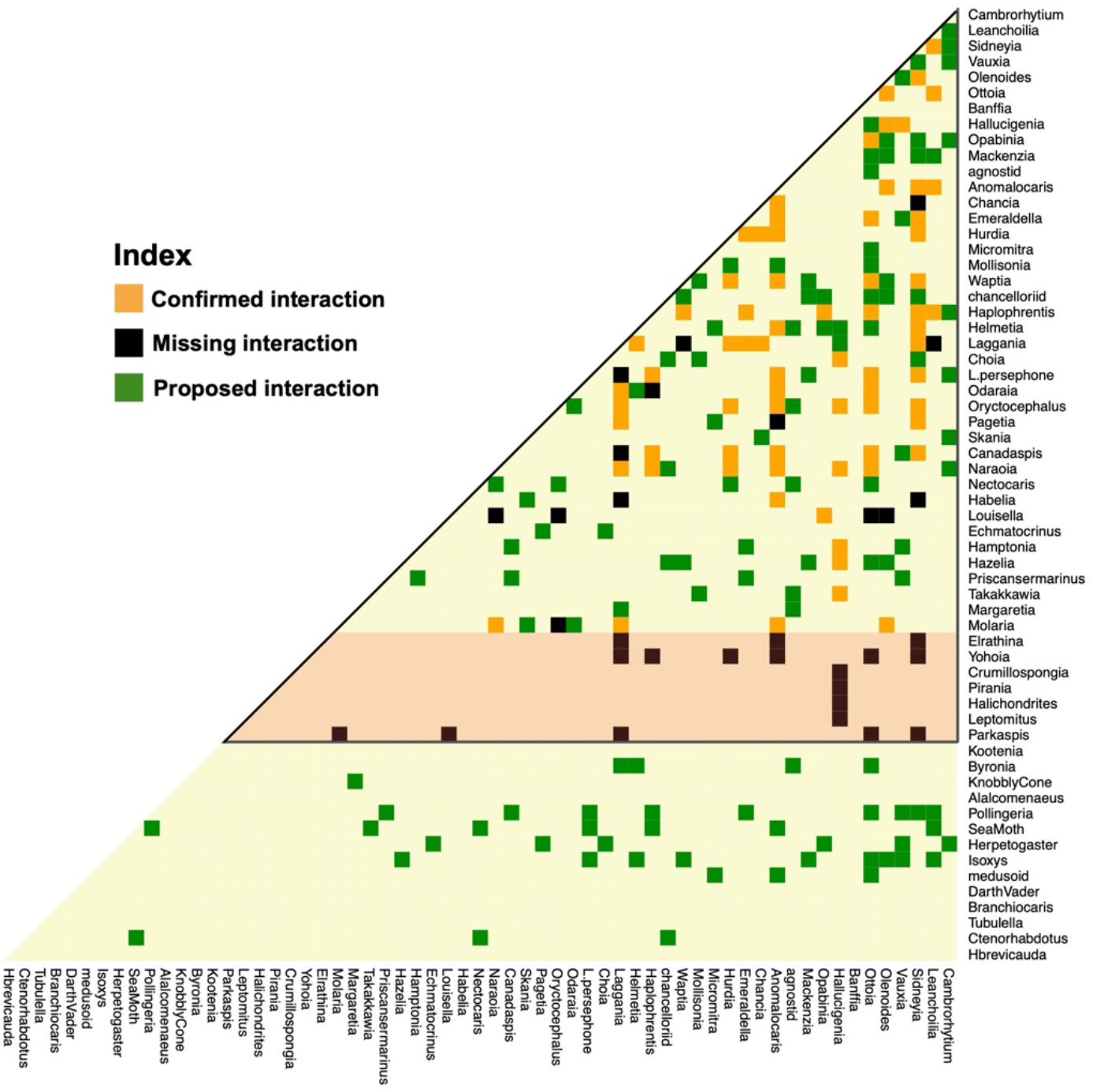
Species interaction half-matrix showing consensus interactions from our analysis, as compared with known trophic interactions from literature (Butterfield., 2000; Dunne et al., 2008; Erwin and Valentine, 2013). Confirmed interactions were proposed in the literature and supported in the correlation analyses here. Missing interactions are reported elsewhere but did not obtain any support from our abundance correlation analyses. Proposed interactions are not currently known from the paleontological literature but are suggested by analyses here. The subset of species interactions inside the black triangle were posited in previous studies (Dunne et al., 2008). The species within the light orange area were numerically rare in our dataset and no statistically robust prediction could be made regarding their interactions. For classification of confirmed interactions, see Figure S8, and for consistency of interaction, see Figure S9.

Detailed ecological analyses of correlation networks may suffer from overestimation problems (Carr et al., 2019; Freilich et al., 2018), but broader-brush categorization of interactions based on abundance correlations can provide novel insights into the functional characteristics of fossil assemblages. Predicted interactions can be supplemented with interactions proposed by paleontological literature, based on gut-contents, morphology or other analyses, to weed out false positives. Other problems raised by earlier studies of paleo-ecological networks (Dunne et al., 2008; Roopnarine, 2010), such as whether correlations capture long term prey-predator and population dynamics, were also explored here. We found that abundance correlation analyses echo results concerning long-term correlations in prey-predator models (Carr et al., 2019) and provide a strong platform for predicting species interactions without reference to prior information concerning the incidence, intensity, or character of those interactions (Figure S4, S5).

Understanding ecological dynamics from fossil data has always been a major challenge, especially for older assemblages. The extraordinary fossil preservation of the Burgess Shale, including the novel Raymond Quarry dataset reported here, provides an exceptional window on possible ecological interactions during an era of major animal evolution. Past studies argued that many properties of modern ecosystem structure first emerged during the Cambrian (Bengston, 2002), and network analyses coupled with proposed trophic interactions highlighted aspects of food-web structure during this period (Dunne et al 2008). When sufficiently strong fossil data are available, analyses of abundance correlation networks, supplemented with models to characterize biases in preservation and interpret species interactions, can reveal unknown or difficult-to-ascertain links in fossilized ecosystems and shed light on trophodynamics over evolutionary time.

## Data and Methods

### Data Collection

The primary data were collected from the Raymond Quarry in the main Burgess Shale site (Figure S1), located ~5 km north of Field, British Columbia, Canada, on a ridge connecting Mt. Field and Wapta Mountain (‘Fossil Ridge’) (figure S1, S2). Other exposures are known (Collins et al. 1983), including outcrops on Mt. Stephen that are lithologically and biostratigraphically equivalent to the Raymond Quarry Member (Fletcher and Collins 1998), such as the outcrops of equivalent strata and fossils across the Kicking Horse Valley on the shoulder of Mt. Stephen (Fletcher and Collins 1998). Consequently, the discussed fauna was not geographically isolated. Moreover, the Raymond Quarry fauna also appears to be autochthonous.

Vertical bedding measurements were determined from an arbitrarily assigned RQ 10.0 m level (equivalent to 21.6 m above the base of C.D. Walcott’s excavations in the Phyllopod Bed). All fossils were labelled according to the bed of occurrence, to the nearest 10cm. Specimens that occurred exactly between two measured levels were assigned to the higher level.

Metadata for each taxon were collected using a literature survey (see references section in ‘metadata_traits.csv’) for the following properties: taxonomic affiliation, habitat, size, motility and preservation potential. Taxonomic affiliation was noted only when there was a majority consensus for the affiliation across sources, otherwise these were omitted from the analysis. Size data were based on the maximum size of specimens found in literature. Motility and habitat were inferred from descriptions of each taxon’s anatomical characteristics. The preservation potential assignments were primarily based on literature descriptions. Hard-bodied taxa are those with biomineralized skeletons, heavily-sclerotized parts, or decay-resistant organic cuticle. Intermediate-group taxa are those with light sclerotization or unsclerotized cuticle. Soft-bodied taxa are those with soft cellular outer layers and soft internal tissues. Enigmatic metazoans (i.e. for which we have no biomineralized/sclerotized preserved parts and have no phylum consensus) were assigned to the soft-bodied group.

### Identifying sub-assemblages

We used ANOSIM (Analysis of Similarity: Clark, 1993; Clark and Warwick, 1994; Bonuso et al., 2002), and SHEBI (S (species richness)-H (information)-E (evenness) analysis for Biofacies Identification: Buzas and Hayek, 1998; Handley et al., 2009) to detect consensus biofacies or sub-assemblages in the sampled region. ANOSIM is a widely used non-parametric, distance-based clustering method. Based on the rank ordering of Bray-Curtis similarities (Clarke and Warwick, 1994), ANOSIM tests for differences in community structure by mixing permutation tests with a general Monte Carlo randomization approach (Hope, 1968; Clarke, 1993). SHEBI recognizes edges between samples of specimens along a (spatial or temporal) transect. It depends upon the anticipated behavior of the evenness metric, which is associated with Shannon-Weaver information, as the number of samples within a single community rises (Handley et al., 2009). Here, each sample was a 10 cm layer of shale with abundance values for each taxon. Each method was conducted on 100 bootstrap simulations of abundance for each 10 cm shale layer. The consensus values from the runs, using both methods, were pooled together, and the mean was used to define the boundaries of the sub-assemblages (named A, B, C and D from oldest to youngest). ANOSIM and SHEBI were implemented using the *vegan* (Okasen et al., 2019) and *foramsv2.0-5* (Aluizio, 2015) packages in R, respectively.

### Agent based models

Agent based models (ABMs) provide an alternative to equation-based simulations for investigating ecological scenarios in a realistic way, along with providing an easy way to incorporate spatial dependence and heterogeneity. Properly implemented, ABMs provide results that match and complement existing ecological theories and experimental evidence (DeAngelis and Grimm, 2014; Karsai et al., 2016). This led to us to choose an ABM implementation over an equation-based implementation, for representing a simple toy model of multiple prey predator interactions. Given that we are dealing with a long term (averaged) ecological abundance dataset of multiple species, there are (i) no equivalent long term complex prey-predator census dynamics of equivalent settings, and (ii) actual prey-predator relations are can only hypothesized based on paleontological evidence (see Dunne et al., 2008), we decided to use a simple ABM implementation that would have as few assumptions as possible and at the same time, also been accepted to give dynamics that have been observed in theory and experiments of prey-predator interactions (Wilensky, 1997, Liebhold et al., 2004; Tobin and Bjørnstad, 2003; Blasius et. al., 2019).

This led us to use the NetLogo language (Wilensky, 1997) and extend a Lotka-Volterra prey-predator (wolf-sheep) base model in the NetLogo library, which replicates simple ecological phenomena among prey and predators (Wilensky, 1997), to create ABMs for purposes of (a) quantifying the impacts of preservation biases, (b) calculating prey-predator correlations, and (c) categorizing interactions.

The primary simulation involved 25 species, with 17 prey and 8 predators, and was based upon an extended implementation of the base Lotka-Volterra wolf-sheep model in the NetLogo library (Wilensky, 1997). All prey fed upon a common resource, which had two parameters attached to it – rate of resource regrowth and initial density of resources (relative to the total area, which was a 500×500 grid). The resource featured an energy content, as did the prey (all prey had equal energy content for simplification). Predators were assigned either a generalist or specialist feeding style. Generalists could eat all types of prey whereas a specialist could only eat one type of prey. Energy was necessary for reproduction, and for simplicity, at each reproduction event we divided the energy between the mother and the offspring, provided that the organism had enough energy to reproduce. The rate of reproduction was controlled as a variable. One million runs of the model were conducted that involved sweeping all parameters throughout their ranges. Each model ran until it reached a stable state (only resource was left, all predators died, or all of the organisms died) or it reached 50,000 time points, whichever occurred first. Data were transferred to R for further processing using the package *RNetLogo* (Thiele, 2017). Each of these models were used as a dataset for (a) testing bias, (b) calculating prey-predator correlations, and (c) categorizing interactions.

The second model was the same as the first/primary model except that each predator species was randomly assigned a different preference for each prey (between 0 and 1). This preference was the probability of a predator consuming a given prey when encountered. One million runs of the model were conducted that involved sweeping all parameters throughout their ranges. Each model ran until it reached a stable state (only resource was left, all predators died, or all of the organisms died) or it reached 50,000 time points, whichever occurred first. Data were transferred to R for further processing using the package *RNetLogo* (Thiele, 2017). Each of these models was then used as a dataset for categorizing interactions.

### Constructing networks

Fossil count data from two adjacent 10 cm layers were combined in each sub-assemblage to increase species coverage for network construction. For each 20cm unit of each sub-assemblage (A through D, excluding A’ and D’ as in Figure 1 on the basis of preservation bias), we iteratively sampled fossils using bootstrap process for 1000 iterations. Using the data for each iteration, we calculated mean correlations between distinct 20 cm blocks for each sub-assemblage across all the bootstrap replicates. Given that some of these interactions can be spurious, we applied partial correlation corrections to the correlation matrices of each sub-assemblage. Next, we performed a Fisher Z-transform of the partial correlation matrices and calculated the probability of observing the estimated Z-scores by chance (based on a normally distributed null distribution). Finally, we used the Benjamini-Hochberg correction of p-values to eliminate those interactions whose corrected correlation p-values exceeded 0.01. This yielded the final set of high-fidelity interactions for each sub-assemblage.

We also created networks on a running time-frame basis where we started at the beginning of the assemblage and repeated the process of network construction as described above for all possible 1.2 m sub-sections created by shifting the analysis frame in 20cm increments.

Pair-wise correlations were calculated for each dataset of the agent-based model simulations and across differently sized time steps as slices for calculating correlations (data were aggregated for each slice) (see Figure S2). We took 100 time-steps as the benchmark for each time slice for constructing networks for all purposes as the correlation saturated at a high enough value and the effects of phase difference and noise were reduced at this time scale of simulations.

### Testing Preservation Bias using Exponential Random Graph Models

Exponential Random Graph Models (ERGMs: Holland and Leinhardt, 1981) are a preferred tool for evaluating how individual variables shape network structures. ERGMs have been used in the past to look at missing data and bias (Robins et al., 2004). We used the *ergm* (Handcock et al., 2019) package in R.

Using the trophic ABM datasets, we assigned each species in each simulated dataset to one of three preservation categories (α, β, or γ) to create a new ‘partially preserved’ dataset whose abundance values were adjusted downward by fixed preservation probabilities where

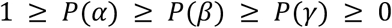

These three categories can be thought of differential preservation categories: for example, in case of body type: hard bodied (preserves well like *α*), intermediate bodied (preserves decently but less than hard body, like *β*) and soft bodied (preserves poorly as compared to other body types; can be denoted by *γ*).

We repeated this procedure 100 times each for all 1 million simulated datasets and then constructed corresponding networks for each of the original and partially preserved datasets. For each of these constructed networks (both altered/partially preserved and original), we calculated the dependence of the network structure on the preservation category (α, β or γ) using ERGMs. We calculated the p-value resulting from the ERGM model for both the altered (partially preserved) networks and the original (intact) networks, as well as their mutual Hamming distance.

The bias coefficient (*B*) is measured in terms of these ERGM p-values of the original and partially preserved networks (*p*_*original*_ and *p*_*partially preserved*_ respectively) as

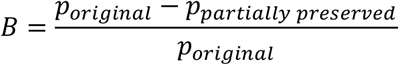

B > 0.5 on this scale corresponds to a change of ~0.2 of Hamming distance (Figure S3). Note that the ABM simulations were used to validate this statistical method before we applied it to the fossil data.

Moreover, this framework can detect biases in network structure based on categorizations, even when the relative preservation potentials among the categories is unknown, but only the categorizations are known. This is a useful property because, although we can assign relative ease of preservation in categorizations of say, body type – assigning the same for habitat and body size might be more complex (see supplementary file ‘metadata_traits.csv’).

Using this framework, we calculated ERGM p-values for each of the networks from the running time-frame analysis, as well as the four sub-assemblages (A-D) of the fossil data. Because we do not have the structure for the unaltered network of the fossil data (i.e. the actual abundance correlation networks from when the burial happened), we assume *p*_*original*_ = 1.0 for these analyses, in order to calculate the bias coefficient. This assumption gives an upper bound on the bias coefficient for our data, as the minimum possible ERGM p-value would be represented by *p*_*original*_ = 1.0 scenario (i.e. no dependence of network structure on any factor), but the actual p-value would usually be lower than this assumed value. As, all reported bias coefficients for the fossil data is based on this assumption, they represent the ‘worst-case’ values.

We performed three sets of analysis in this regard: body type, body size, and habitat affiliation. In each set, there were three categories, which were pre-determined (see supplementary file ‘metadata_traits.csv’ for details), according to available paleontological evidence. The body type categorizations were based on preservation or fossilization potential, as described in ‘Data collection’ sub-section of Methods – namely, hard bodied, soft bodied and intermediate, based on literature descriptions. Body size categorizations were <15 cm, 15 - 30 cm, > 30 cm maximum size. Habitat types were categorized into endobenthic/epibenthic, nektobenthic, nektonic/pelagic, based on literature survey (see references section in ‘metadata_traits.csv’).

### Categorization of interactions and comparison with ABM

We subjected the distributions of abundance correlations to a maximum likelihood analysis to identify appropriate Gaussian basis functions. The mean, and standard deviation for a pre-defined number of Gaussian basis functions were determined using a general simplex-based optimization algorithm (Nelder and Mead 1965). We compared results assuming 1 through 6 possible Gaussian basis functions and identified the optimal number of such functions for the correlation data using the likelihood-ratio test. All the scripts were implemented in R.

Once the number and nature of the Gaussians were estimated (for sub-assemblages A-D and for the running time-frame analysis), we used pairwise Kolmogorov-Smirnov (KS) tests between basis Gaussians to obtain a pairwise similarity matrix. The number of clusters of Gaussian basis functions was determined using spectral analysis based on the pairwise similarity matrix obtained from KS analysis and the gap statistic (Tibshirani et al., 2000). An advantage of using the gap statistic is that it does not pre-assume the number of required clusters (Tibshirani et al., 2000). All scripts were implemented in R using the package *cluster*. Four zones of clustering were determined using this method and are termed categories of interactions. Each category is shown in Figure 2 (a) using the range of all associated Gaussian means.

Next, from the ABM datasets, we identified the nature (i.e., generalist-prey, specialist-prey, competition, apparent competition) of all interactions whose partial correlations fell within ranges of the Gaussian mean clusters and plotted their relative occurrences in Figure 2c.

Consensus interactions were calculated from the running time-frame analysis and were defined as the high-fidelity interactions (or statistically corrected correlations) that stayed in the same interaction category for more than 50% of the time it occurred for a given pair of taxa. These results have been plotted in figures 3, S8 and S9.

### Stochastic Block Models (SBMs) and Equitability analysis

The stochastic block model (SBM) is a tool, used to detect community structure in a network, where communities can be defined as multi-node subcomponents (or, blocks) of the network in which edges are more common within than between communities (Karrer and Newman, 2011).

We applied the framework of SBMs to the networks constructed from fossil data to understand the associations among species. In particular, we sought to understand whether the correlations on which those networks were built represented shared motility or habitat variables rather than species interactions. At each network level (which were calculated at a sub-assemblage level A-D and also at a fine time scale level), we used integrated classification likelihood (ICL) to calculate the number of clusters/blocks. On each block/cluster, we annotated the taxa in those blocks using the metadata on habitat and motility (separately; see supplementary file ‘metadata_traits.csv’ for details) and used the *mixer* package (Latouche et al., 2012) to find the distribution of annotated categories across blocks at a given network level. In order to numerically represent it, we calculated Shannon’s equitability index (SEI), which is the normalized Shannon’s diversity coefficient, based on the categories of habitat/motility for each block/cluster – and to estimate a network level (for a sub-assemblage/fine time scale analysis) average – we found the weighted average (on basis on number of taxa in each block/cluster) of SEI over all the blocks/clusters in the given network. We then plotted this average network SEI value at the fine scale analysis level in Figure S10(c).

SEI points at the dominance of a specific type of say, habitat/motility, on the taxa involved in interactions within a calculated cluster. If a given cluster is highly dominated by a single type of habitat/motility, SEI would be very low and would be 0 if it is only type. As SEI is normalized diversity index, the highest possible value of 1 occurs when all the types are equally probable. This is not the case with our fossil data and hence, we calculated the maximum empirical value possible with the data (Figure S10(c)) for both habitat and motility separately. In addition, to make sense of how biased the correlational values are, we calculated the SEIs, for both habitat and motility, where a dominant type is equal to 95% of a given cluster and others are equally distributed in the remaining fraction (Figure S10(c)).

## Data availability

All relevant data needed to recreate the results are provided as supplementary material. Abundance values for all the taxa at 10 cm resolution in RQ can be found in ‘abundance_data.csv’; ecological habits, taxonomic affinity, body type categorization and size metadata for all the taxa can be found in ‘metadata_traits.csv’. Relevant final data are also provided for reference – ‘trophic_interaction_matrix.csv’ contains the consensus trophic interactions, and ‘competitive_interaction_matrix.csv’ contains the consensus competitive/negative interactions. Simplified ABM simulation and network analysis code can be found at: github.com/anshuman21111/cambrian-fossil-networks.

## Acknowledgements

We thank Nicholas J Butterfield, Phillip Staniczenko, Jennifer Dunne, Michelle Girvan, Hector Corrada-Bravo, Tracy Chen, Sushant Patkar, Roozbeh Bassirian, Doug Erwin, Philip Johnson and Jake Weissman for their helpful suggestions and discussions. We acknowledge help from Jack O Shaw in collecting some of the metadata for our analysis. A.S.’s contribution to this research was supported in part by NSF award DGE-1632976.

## Supplementary Materials

**Figure S1:**
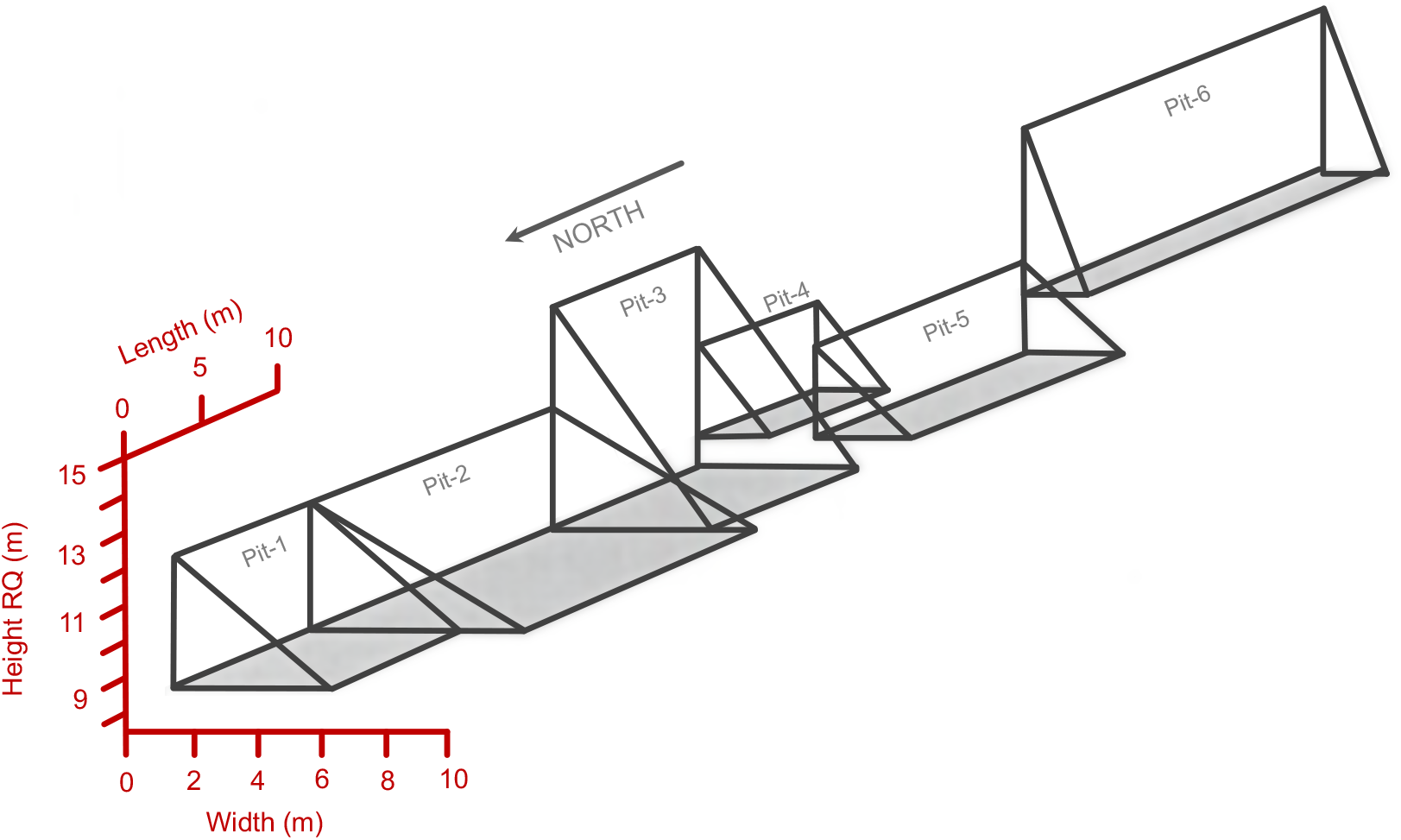
Schematic of excavations in the Raymond Quarry Member. The original Raymond Quarry was located within Pit #5. The northernmost extent of Pit # l is 23 m south of the Cathedral Escarpment.

**Figure S2:**
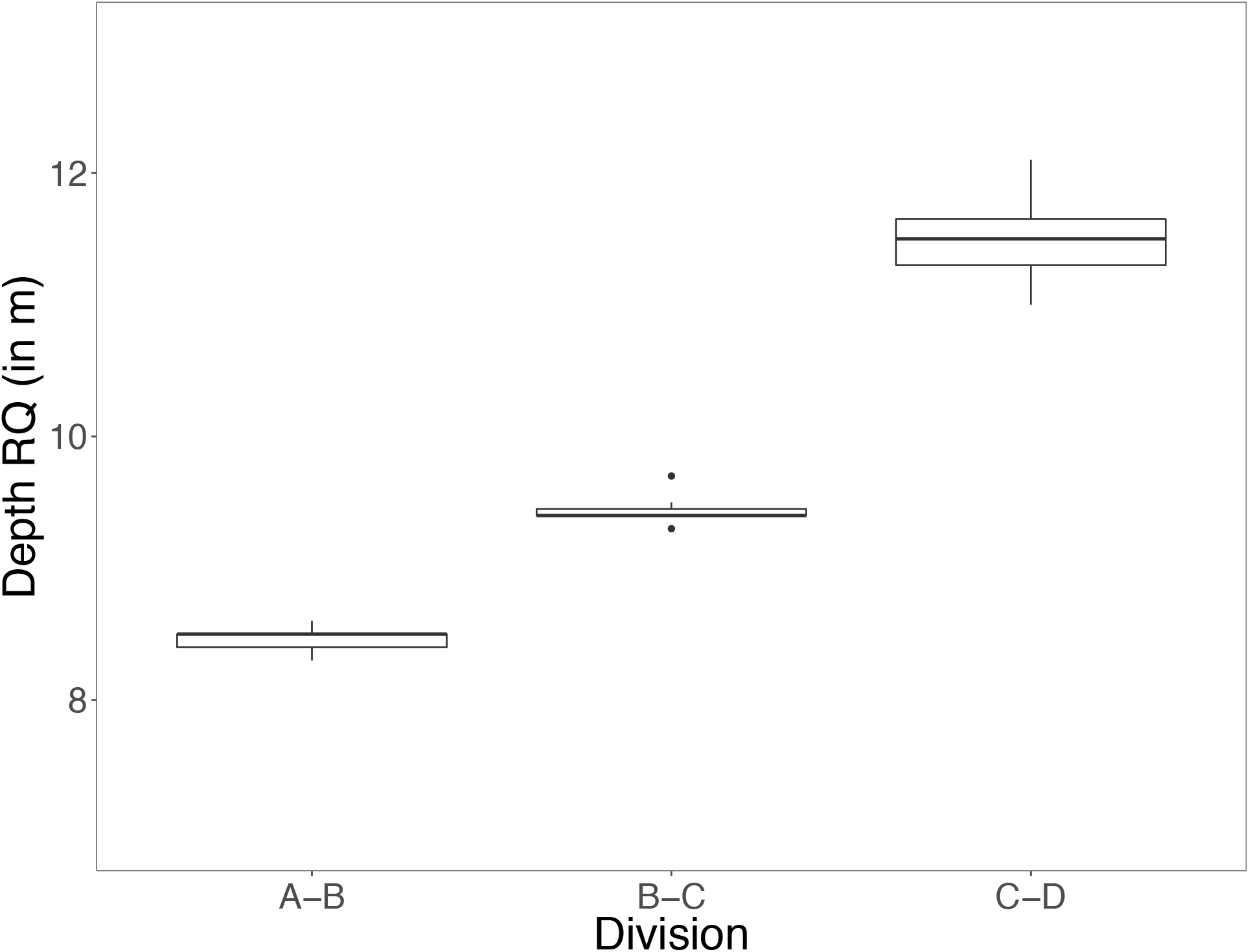
Biofacies detected using consensus of statistical methods ANOSIM and SHEBI for the boundaries between the Raymond Quarry sub-assemblages A-B, B-C, and C-D.

**Figure S3:**
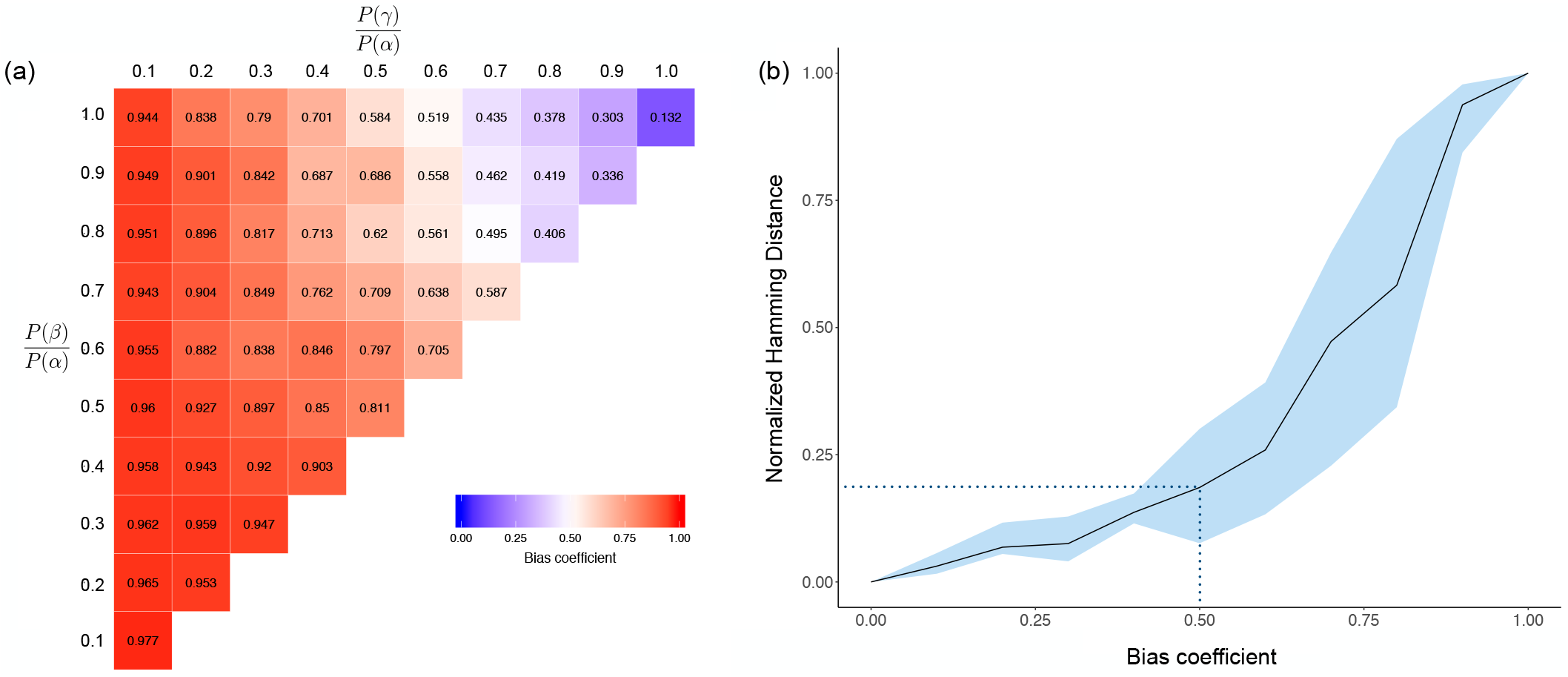
Preservation Bias: (a) Plot of bias coefficient as measured from the agent-based model for three categories of preservation probability: α, β and γ; (b) Differences between networks quantified using the Hamming distance (larger values imply greater deviation) as a function of the bias coefficient. The plot compares unaltered ABM networks with ABM networks whose structure was obscured by preservation biases; standard errors are calculated across alternative ABM simulations. Note the nonlinear dependence of Hamming distance on bias that increases steeply beyond bias coefficient of ~0.50.

**Figure S4:**
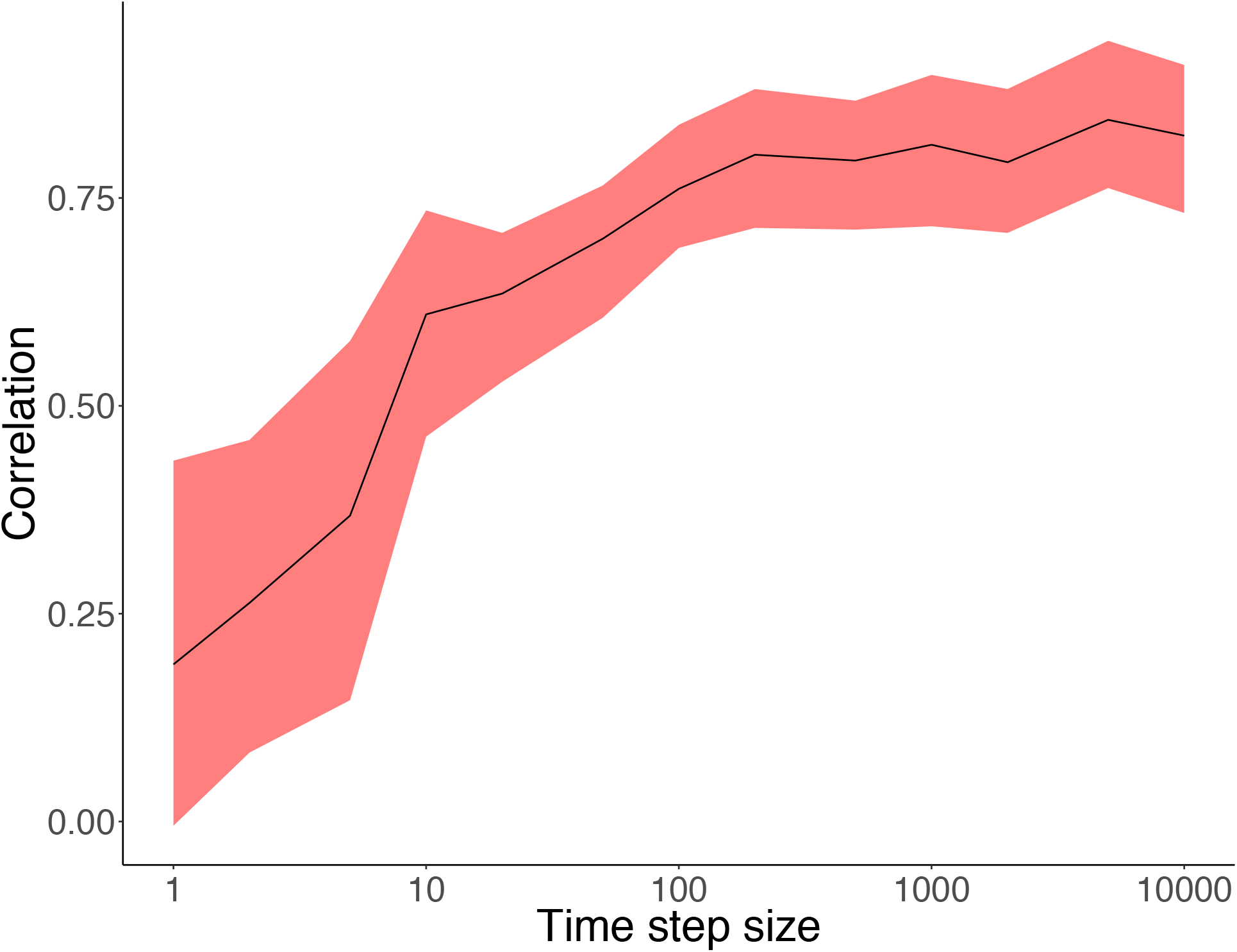
Correlation between a prey and its specialist predator in ABM output based on 10 sampled population sizes separated by the specified time steps. The cloud of standard error values was calculated using all the ABM runs with different initial conditions. Each ABM simulation was taken as a separate dataset and the correlation was calculated for each pair of prey and specialist predators in all the datasets. Increasing the number of sampled counts used to calculate the correlation substantially decreases the width of the error cloud.

**Figure S5:**
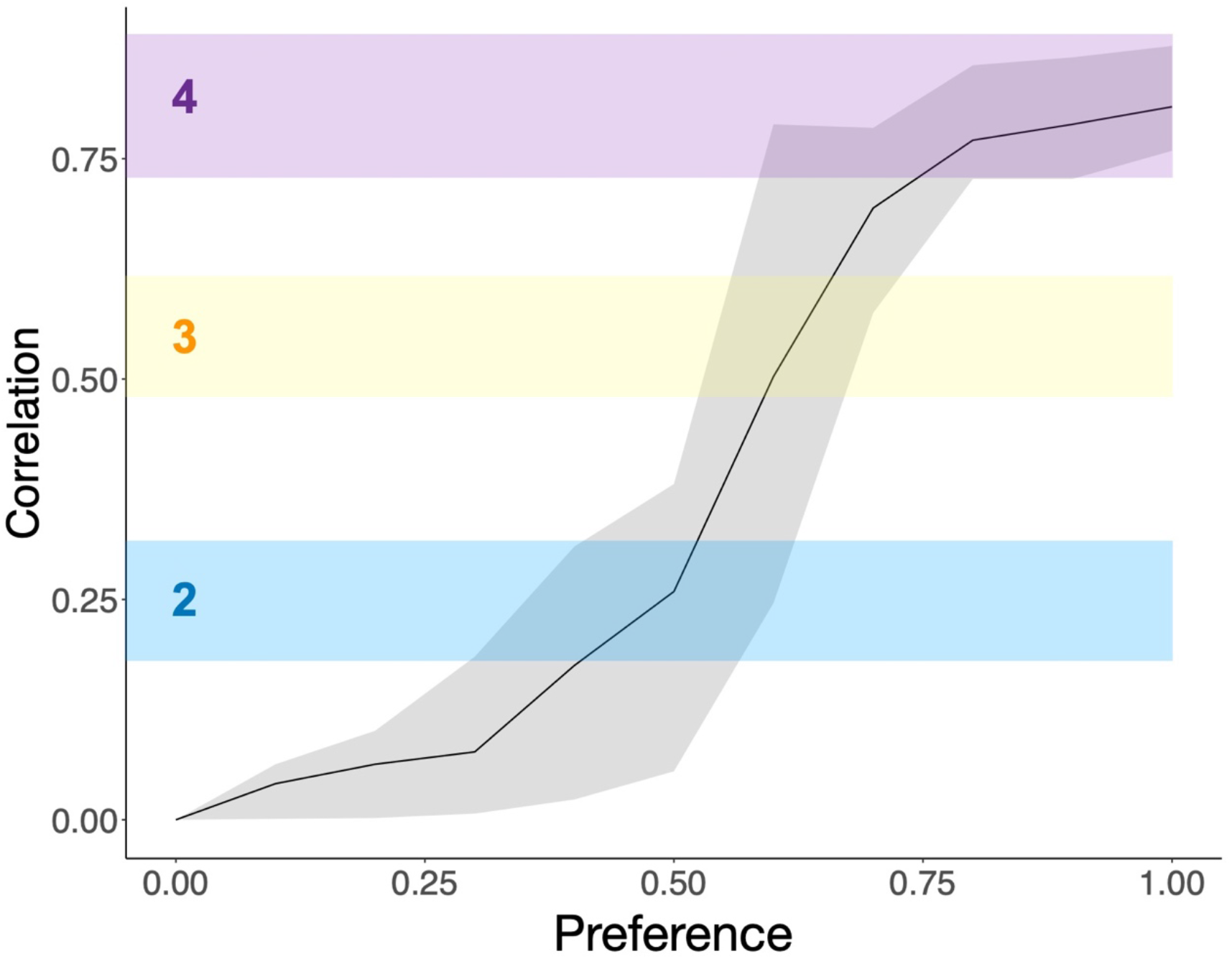
Correlation change and preference/affinity for prey in an ABM with standard error (run for 10,000 different ABM scenarios). Mapped onto the correlation axis are the three trophic interaction categories 2, 3, and 4 from the spectral analysis.

**Figure S6:**
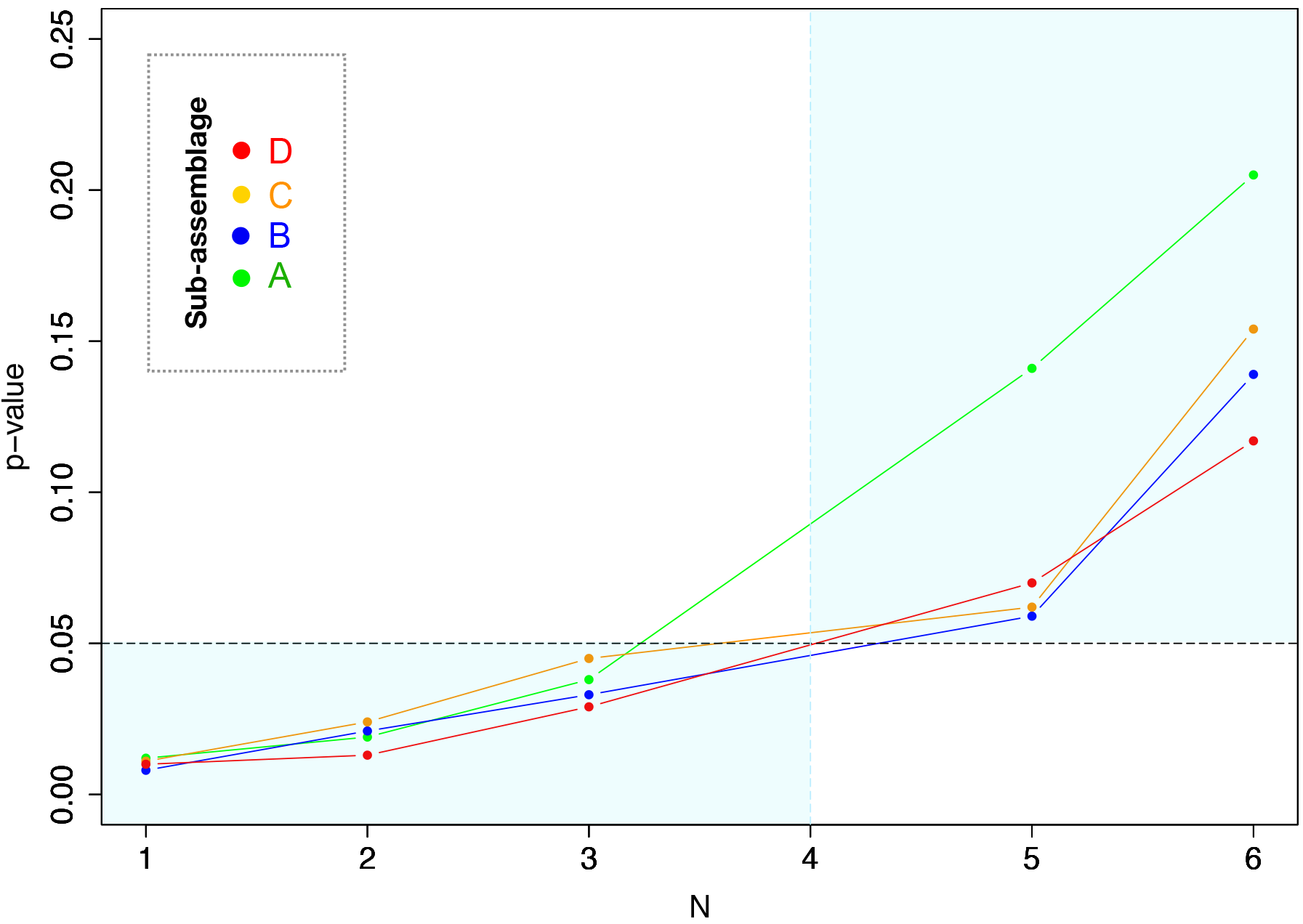
p-values from chi-square test of likelihood ratio tests for nested models that differ in the number of Gaussians (N) as compared with a model with 4 Gaussians (for sub-assemblages A-D). The areas highlighted in light blue signify where the model with four Gaussians performs significantly better. All the points for all sub-assemblages lie in the region implying that a model with four basis Gaussians is the best model to pick.

**Figure S7:**
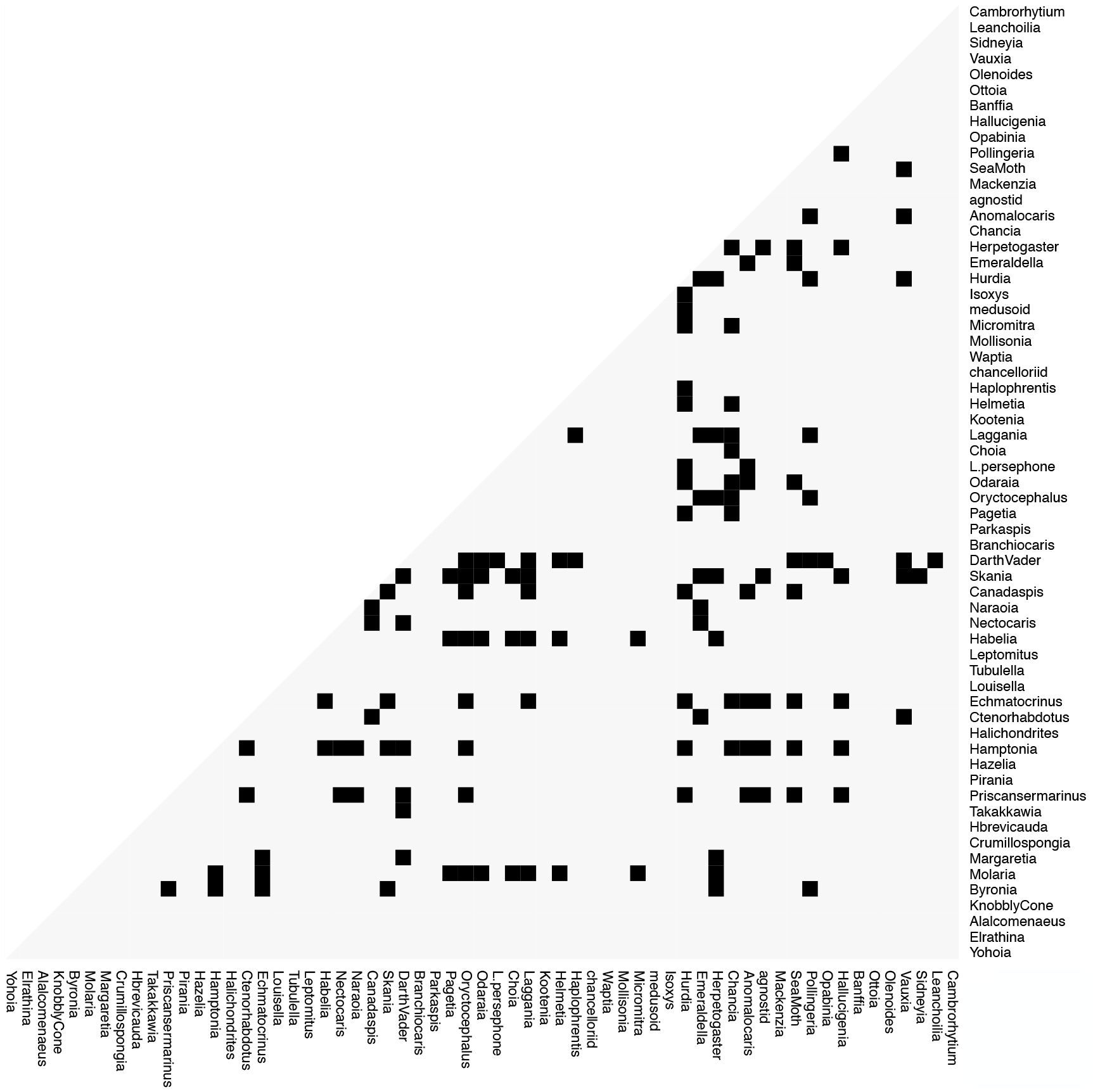
Species interaction half-matrix representing competitive interactions among the fossil taxa proposed on the basis of abundance correlation network analyses. As it is more difficult to test competitive interactions through anatomical or other paleo-ecological evidence, we did not pursue these further, instead focused only on trophic interactions.

**Figure S8:**
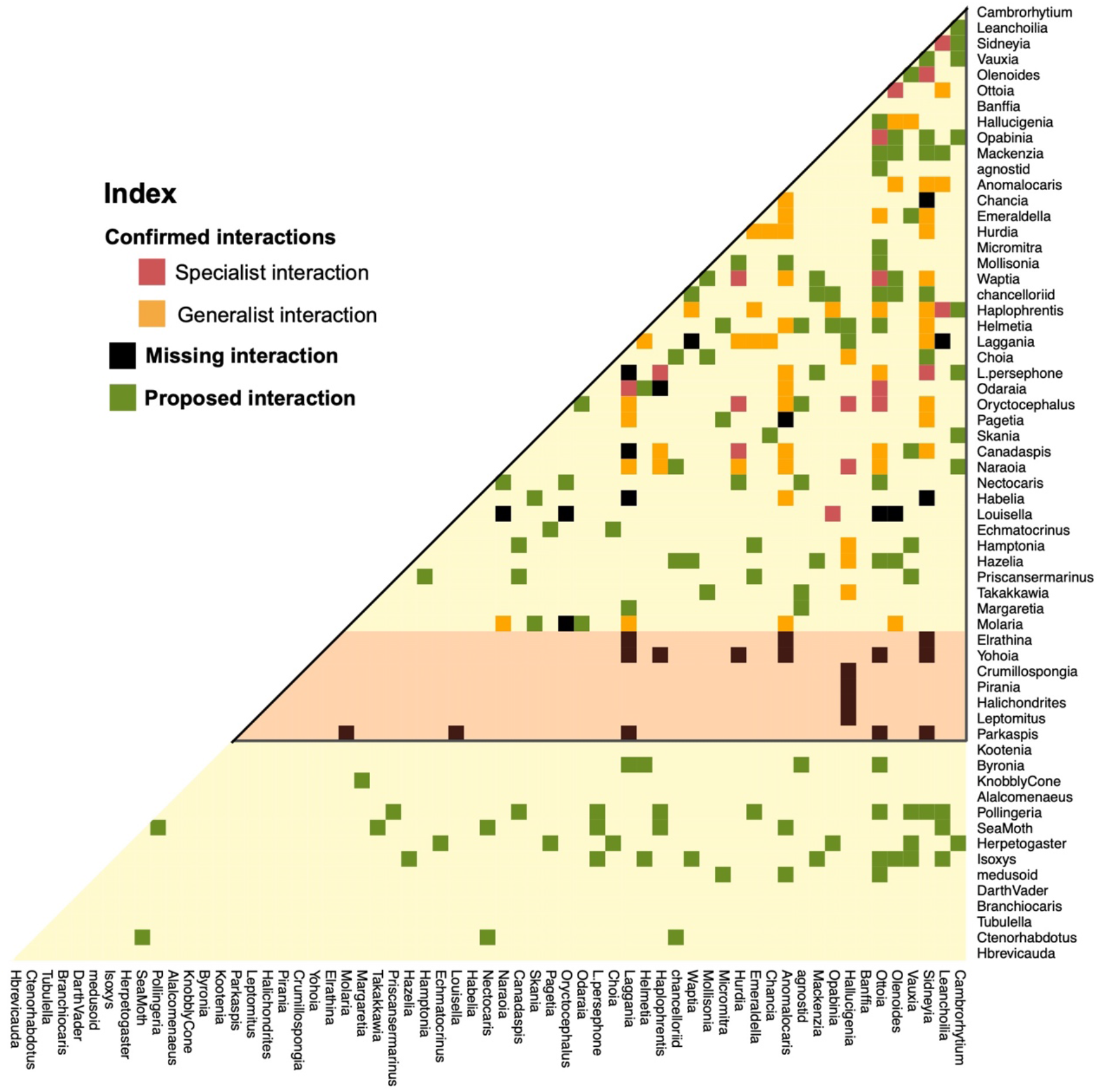
Species interaction matrix showing the results from the categorization analysis, as compared with known trophic interactions from literature (Butterfield., 2000; Dunne et al., 2008; Erwin and Valentine, 2013), along with breakdown of the confirmed trophic interactions into specialist and generalist categories. Confirmed interactions were proposed in the literature and supported in the correlation analyses here. Missing interactions are reported elsewhere but did not obtain any support from our abundance correlation analyses. Proposed interactions are not currently known from the paleontological literature but are suggested by analyses here. The subset of species interactions within the black triangle are known from previous studies (Dunne et al., 2008). The species within the light orange area were numerically rare in our dataset and no statistically robust prediction could be made regarding their interactions.

**Figure S9:**
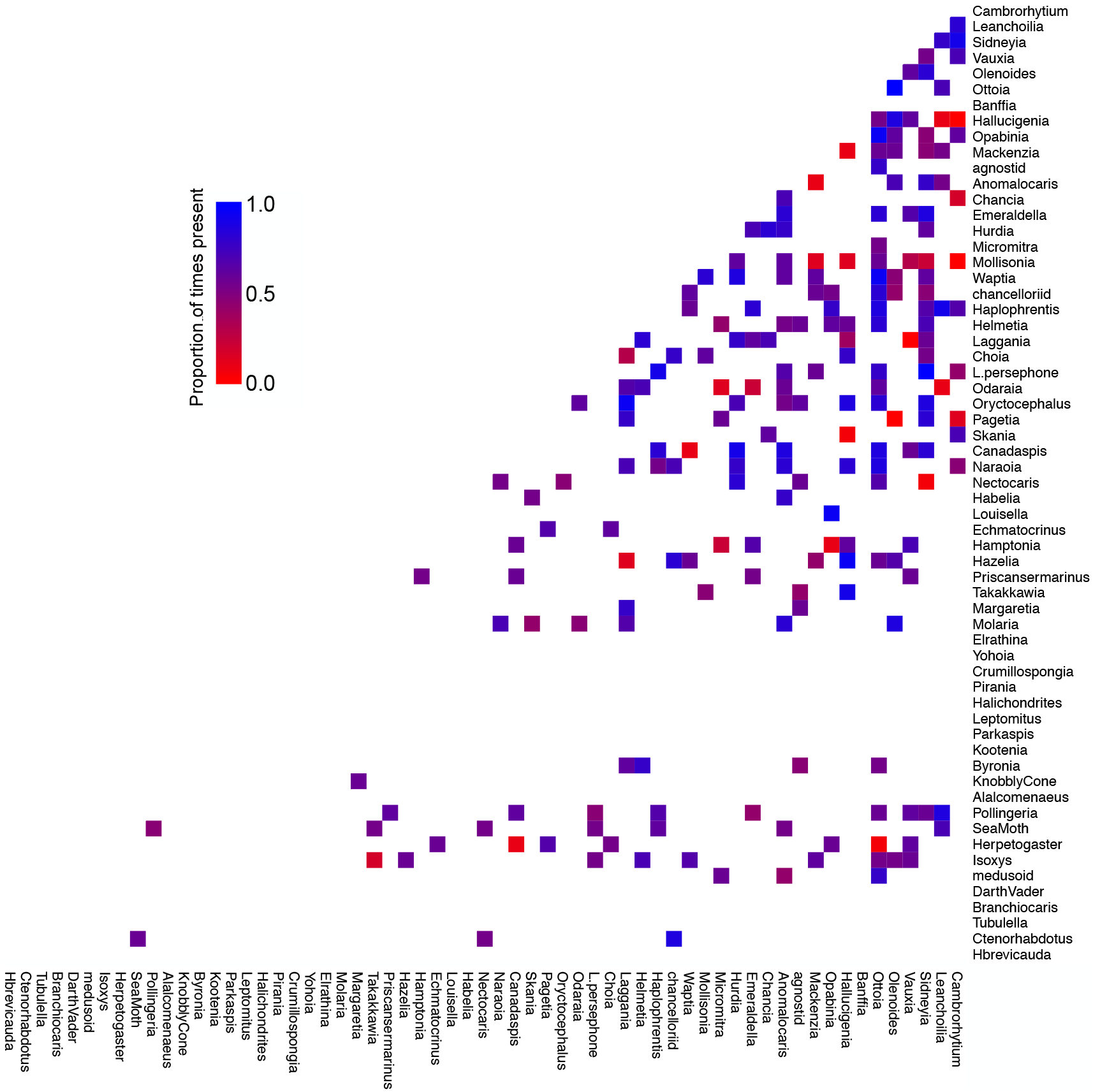
Heatmap showing the proportion of times high fidelity interactions were observed in the same categorization. 82.37% of the high-fidelity interactions are present consistently in the same interaction category more than 50% of the time (of co-occurrences of the pair of species) and are termed as consensus interactions. These consensus interactions are presented in the species interaction half-matrix in figure 3 and S8.

**Figure S10:**
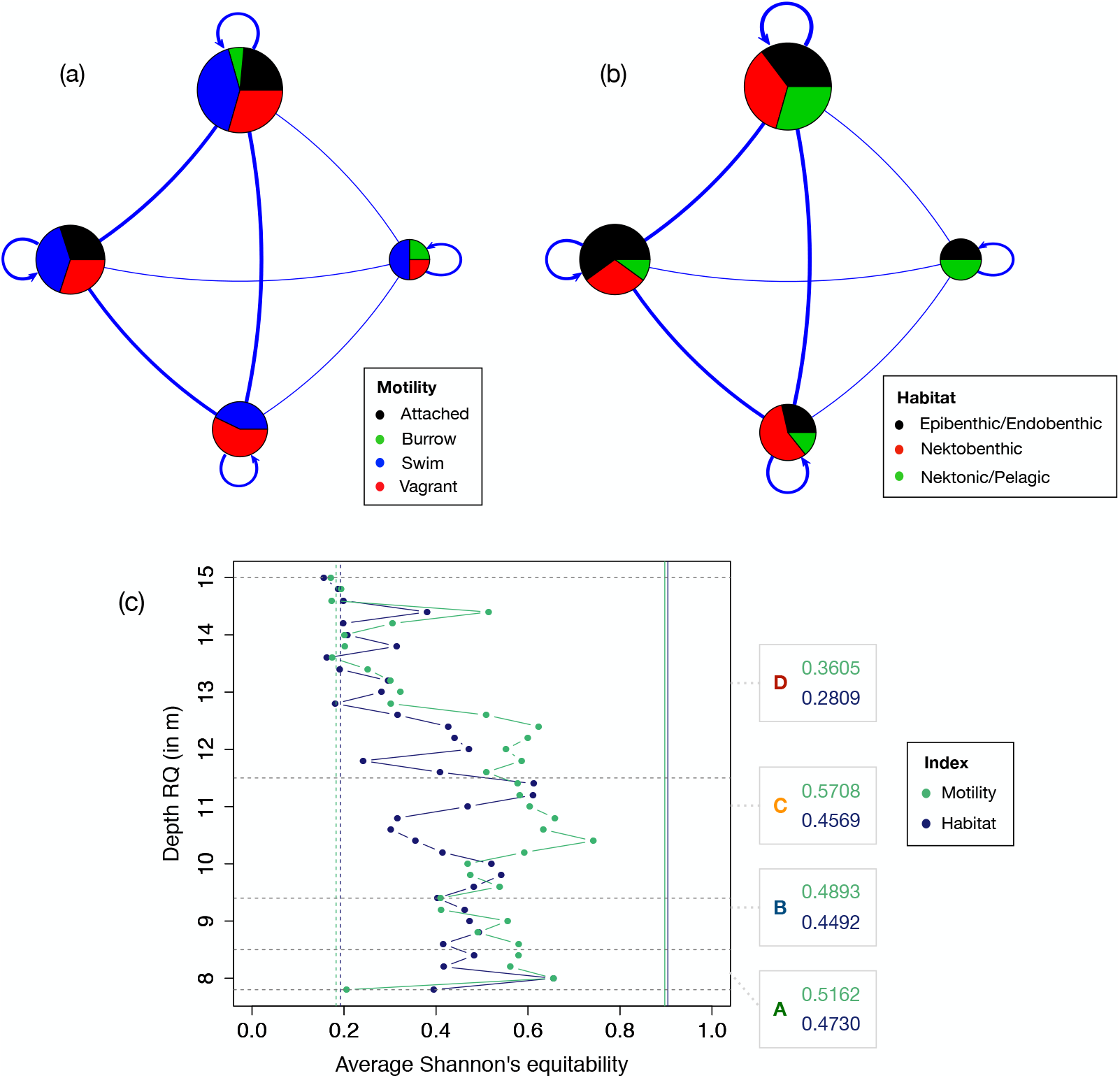
A stochastic block model of the overall network (sub-assemblages A-D) is shown here for representation purposes, which depicts four distinct blocks of taxa, estimated using integrated classification likelihood, that interact more within blocks than across blocks for motility classes (a) and habitat affinity classes (b). The lines between groups and their width denotes interaction between two blocks of taxa. Please note that the width of the self-loop here is not meaningful, other than the fact that the taxa within a block interact among themselves. The size of the block refers to the number of taxa in that block. The model has been overlaid here with (a) motility data, and (b) habitat data. Clearly, organisms belonging to particular motility or habitat classes do not fall consistently into SBM blocks. This, in turn, suggests that habitat/motility similarity does not transform into very strong associations through correlations. Please note that the number of blocks varied in different sub-assemblages and running time series analysis, as it is dependent on the structure of the network. In (c) we use this SBM model to estimate average Shannon’s equitability index (SEI) (methods) for the running timeframe analysis and for sub-assemblages (A-D). The grey horizontal dotted lines demarcate the sub-assemblages (A-D). The solid lines (separate for motility and habitat) depict the maximum theoretical value of SEI possible using our data and the dashed lines represent a theoretical SEI for 95% dominance by a single type of habitat/motility (Methods). One can see that, the SEI values depict no dependence of correlation network community structure on habitat or motility, except the start and end of the assemblage – where a weak dependence cannot be ruled out. This is in lines with the bias coefficient (methods, Figure 1(b)).

## References

1. Aluizio R. (2015). forams: Foraminifera and Community Ecology Analyses. R package version 2.0-5. https://CRAN.R-project.org/package=forams

2. Armitage, D. W., & Jones, S. E. (2019). How sample heterogeneity can obscure the signal of microbial interactions. The ISME journal, 13(11), 2639–2646.

3. Barberán, A., Bates, S. T., Casamayor, E. O., & Fierer, N. (2012). Using network analysis to explore co-occurrence patterns in soil microbial communities. The ISME journal, 6(2), 343.

4. Bascompte J (2010), “Structure and Dynamics of Ecological Networks”, Science 329:5993, pp. 765–766

5. Bengtson, S. (2002). Origins and early evolution of predation. The Paleontological Society Papers, 8, 289–318.

6. Blasius, B., Rudolf, L., Weithoff, G., Gaedke, U. and Fussmann, G.F., 2019. Long-term cyclic persistence in an experimental predator–prey system. Nature, pp.1–5.

7. Butterfield, N. J. (2003). Exceptional fossil preservation and the Cambrian explosion. Integrative and Comparative Biology, 43(1), 166–177.

8. Buzas, M. A., & Hayek, L. A. C., 1998. SHE analysis for biofacies identification. Journal of Foraminiferal Research.

9. Carbone, C., & Narbonne, G. M. (2014). When Life Got Smart: The Evolution of Behavioral Complexity Through the Ediacaran and Early Cambrian of NW Canada. Journal of Paleontology, 88(02), 309–330.

10. Carr, A., Diener, C., Baliga, N. S., & Gibbons, S. M. (2019). Use and abuse of correlation analyses in microbial ecology. The ISME journal, 13(11), 2647–2655.

11. Clarke, K. R., & Warwick, R. M. (1994). Similarity-based testing for community pattern: the two-way layout with no replication. Marine Biology, 118(1), 167–176.

12. Caron, J. B., Gaines, R. R., Aria, C., Mángano, M. G., & Streng, M. (2014). A new phyllopod bed-like assemblage from the Burgess Shale of the Canadian Rockies. Nature Communications, 5, 3210.

13. Colles, A., Liow, L. H., & Prinzing, A. (2009). Are specialists at risk under environmental change? Neoecological, paleoecological and phylogenetic approaches. Ecology letters, 12(8), 849–863.

14. Conway Morris S (1986) The community structure of the Middle Cambrian Phyllopod Bed (Burgess Shale). Palaeontology 29: 423–467.

15. Conway Morris, S. (1989). Burgess Shale faunas and the Cambrian explosion. Science, 246(4928), 339–346.

16. Delmas E, Besson M, Brice M H, Burkle L A, Dalla Riva G V, Fortin M J, Gravel D, Guimarães Jr P R, Hembry D H, Newman E A, Olesen J M (2017), “Analysing ecological networks of species interactions” Biological Reviews

17. Devereux, M. G. (2001). Palaeoecology of the Middle Cambrian Raymond Quarry Fauna, Burgess Shale, British Columbia.

18. Dormann, C. F., Fründ, J., & Schaefer, H. M. (2017). Identifying causes of patterns in ecological networks: opportunities and limitations. Annual Review of Ecology, Evolution, and Systematics, 48, 559–584.

19. Dunne J A, Williams R J, Martinez N D, Wood R A & Erwin D H (2008) “Compilation and network analyses of Cambrian food webs” PLoS Biology 6(4), e102

20. Dunne J A, Labandeira C C and Williams R J (2014) “Highly resolved early Eocene food webs show development of modern trophic structure after the end-Cretaceous extinction” Proceedings of the Royal Society of London B: Biological Sciences 281(1782), 20133280.

21. Erwin, D. H., Laflamme, M., Tweedt, S. M., Sperling, E. A., Pisani, D., & Peterson, K. J. (2011). The Cambrian conundrum: early divergence and later ecological success in the early history of animals. Science, 334(6059), 1091–1097.-1097.

22. Erwin, D. H., & Valentine, J. W. (2012). The Cambrian explosion: the construction of animal biodiversity. Roberts.

23. Flannery Sutherland, J. T., Moon, B. C., Stubbs, T. L., & Benton, M. J. (2019). Does exceptional preservation distort our view of disparity in the fossil record? Proceedings of the Royal Society B, 286(1897), 20190091.

24. Freilich, M.A., Wieters, E., Broitman, B.R., Marquet, P.A. and Navarrete, S.A., (2018). Species co‐occurrence networks: Can they reveal trophic and non‐trophic interactions in ecological communities?. Ecology, 99(3), pp.690–699.

25. Handcock, M., Hunter, D., Butts, C., Goodreau, S., Krivitsky, P., and Morris, M. (2019). _ergm: Fit, Simulate and Diagnose Exponential-Family Models for Networks_. The Statnet Project (https://statnet.org). R package version 3.10.4. https://CRAN.R-project.org/package=ergm.

26. Ings, T.C., Montoya, J.M., Bascompte, J., Blüthgen, N., Brown, L., Dormann, C.F., Edwards, F., Figueroa, D., Jacob, U., Jones, J.I. and Lauridsen, R.B., 2009. Ecological networks–beyond food webs. Journal of Animal Ecology, 78(1), pp.253–269.

27. Jordano, P., (2016). Sampling networks of ecological interactions. Functional Ecology, 30(12), pp.1883–1893.

28. Karrer, Brian, and Mark EJ Newman. “Stochastic blockmodels and community structure in networks.” Physical review E 83, no. 1 (2011): 016107.

29. Kidwell, S. M., Bosence, D. W., Allison, P. A., & Briggs, D. E. G., 1991. Taphonomy and time-averaging of marine shelly faunas. Taphonomy: releasing the data locked in the fossil record. Plenum, New York, 115–209.

30. Koch C (1978), “Bias in the published fossil record” Paleobiology,4:3, pp. 367–372

31. Latouche, P., Birmelé, E. and Ambroise, C. (2012), Variational Bayesian inference and complexity control for stochastic block models. Statistical Modelling, SAGE Publications, 12, 1, 93–115.

32. Lau M K, Borrett S R, Baiser B, Gotelli N J, and Ellison A M (2017), “Ecological network metrics: opportunities for synthesis”, Ecosphere 8:8, e01900

33. Lee, Michael SY, Julien Soubrier, and Gregory D. Edgecombe. “Rates of phenotypic and genomic evolution during the Cambrian explosion.” (2013). Current Biology 23, no. 19: 1889–1895.

34. Liebhold, A., Koenig, W. D., & Bjørnstad, O. N. (2004). Spatial synchrony in population dynamics. Annu. Rev. Ecol. Evol. Syst., 35, 467–490.

35. Mángano, M. G., & Buatois, L. A. (2014). Decoupling of body-plan diversification and ecological structuring during the Ediacaran–Cambrian transition: evolutionary and geobiological feedbacks. Proceedings of the Royal Society B: Biological Sciences, 281(1780), 20140038.

36. Martin, R.E., 1999. Taphonomy: a process approach (Vol. 4). Cambridge University Press.

37. Mitchell, E.G., Harris, S., Kenchington, C.G., Vixseboxse, P., Roberts, L., Clark, C., Dennis, A., Liu, A.G. and Wilby, P.R., 2019. The importance of neutral over niche processes in structuring Ediacaran early animal communities. Ecology letters, 22(12), pp.2028–2038.

38. Mitchell, E.G. and Butterfield, N.J., “Spatial analyses of Ediacaran communities at Mistaken Point”. Paleobiology (2018), 44(1), pp.40–57.

39. Mitchell, J. S. (2015). Preservation is predictable: quantifying the effect of taphonomic biases on ecological disparity in birds. Paleobiology, 41(2), 353–367.

40. Muscente, A.D., Bykova, N., Boag, T.H., Buatois, L.A., Mángano, M.G., Eleish, A., Prabhu, A., Pan, F., Meyer, M.B., Schiffbauer, J.D. and Fox, P., “Ediacaran biozones identified with network analysis provide evidence for pulsed extinctions of early complex life”. Nature communications (2019): 10(1), pp.1–15.

41. Muscente A D, Prabhu A, Zhong H, Eleish A, Meyer M B, Fox P, Hazen R M, and Knoll A H. “Quantifying ecological impacts of mass extinctions with network analysis of fossil communities.” Proceedings of the National Academy of Sciences (2018): 201719976.

42. Nelder, J. A. and Mead, R. (1965). A simplex algorithm for function minimization. Computer Journal, 7, 308–313

43. Oksanen, J., Blanchet, F.G., Friendly, M., Kindt, R., Legendre, P., McGlinn, D., Minchin, P.R., O’Hara, R.B., Simpson, G.L., Solymos, P., Stevens, M.H.H., Szoecs, E. and Wagner, H. (2019). vegan: Community Ecology Package. R package version 2.5-6. https://CRAN.R-project.org/package=vegan

44. Olson, E.C., Behrensmeyer, A.K. and Hill, A.P., 1980. Taphonomy: its history and role in community evolution. Fossils in the making: vertebrate taphonomy and paleoecology, pp.5–19.

45. Orr, P. J., Benton, M. J., & Briggs, D. E. (2003). Post-Cambrian closure of the deep-water slope-basin taphonomic window. Geology, 31(9), 769–772.

46. Paine R T (1969). “The Pisaster-Tegula interaction: prey patches, predator food preference, and intertidal community structure”, Ecology 50, pp. 950–961

47. Piechnik, D.A., Lawler, S.P. and Martinez, N.D., 2008. Food‐web assembly during a classic biogeographic study: species’ “trophic breadth” corresponds to colonization order. Oikos, 117(5), pp.665–674.

48. Poisot T, Stouffer D B, and Kefi S (2016), “Describe, understand and predict: Why do we need networks in ecology?” Functional Ecology 30, pp. 1878–1882

49. Robins, G., Pattison, P., & Woolcock, J. (2004). Missing data in networks: exponential random graph (p∗) models for networks with non-respondents. Social Networks, 26(3), 257–283.

50. Roopnarine P D, Angielczyk K D, Wang S C and Hertog R (2007) “Trophic network models explain instability of Early Triassic terrestrial communities”, Proceedings of the Royal Society of London B: Biological Sciences, 274(1622), pp. 2077–2086.

51. Roopnarine, P. D. (2010). Networks, extinction and paleocommunity food webs. The Paleontological Society Papers, 16, 143–161.

52. Ryall, K. L., & Fahrig, L. (2006). Response of predators to loss and fragmentation of prey habitat: a review of theory. Ecology, 87(5), 1086–1093.

53. Saleh, F., Antcliffe, J.B., Lefebvre, B., Pittet, B., Laibl, L., Peris, F.P., Lustri, L., Gueriau, P. and Daley, A.C., 2020. Taphonomic bias in exceptionally preserved biotas. Earth and Planetary Science Letters, 529, p.115873.

54. Seilacher, A., Buatois, L. A., & Mángano, M. G. (2005). Trace fossils in the Ediacaran–Cambrian transition: behavioral diversification, ecological turnover and environmental shift. Palaeogeography, Palaeoclimatology, Palaeoecology, 227(4), 323–356.

55. Smith A B (2001), “Large–scale heterogeneity of the fossil record: implications for Phanerozoic biodiversity studies”, Proceedings of the Royal Society of London B: Biological Sciences, 365, pp. 351–367

56. Starrfelt, J., & Liow, L. H. (2016). How many dinosaur species were there? Fossil bias and true richness estimated using a Poisson sampling model. Philosophical Transactions of the Royal Society B: Biological Sciences, 371(1691), 20150219.

57. Thiele J.C. (2017). RNetLogo: Provides an Interface to the Agent-Based Modelling Platform ‘NetLogo’. R package v.1.0-4. https://cran.r-project.org/package=RNetLogo

58. Tobin, P. C., & Bjørnstad, O. N. (2003). Spatial dynamics and cross‐correlation in a transient predator–prey system. Journal of Animal Ecology, 72(3), 460–467.

59. Tibshirani, R., Walther, G., & Hastie, T. (2001). Estimating the number of clusters in a data set via the gap statistic. Journal of the Royal Statistical Society: Series B (Statistical Methodology), 63(2), 411–423.

60. Vannier, J., & Chen, J. (2005). Early Cambrian food chain: new evidence from fossil aggregates in the Maotianshan Shale biota, SW China. Palaios, 20(1), 3–26.

61. Zhang, W. (2011). Constructing ecological interaction networks by correlation analysis: hints from community sampling. Network Biology, 1(2), 81.

